# Dispersal, maturation and recruitment in a long-lived, trans-continentally migrating bird

**DOI:** 10.1101/2024.05.17.594760

**Authors:** Paweł Mirski, Wouter Vansteelant, Patrik Byholm

## Abstract

Natal dispersal is a multi-step process. It commences when a juvenile departs from its natal site and concludes when it settles to breed for the first time. During this interim period, dispersing individuals of long-lived species undergo a wandering phase, which may span years. This phase remains one of the least studied aspects of species’ life history. We utilized a unique GPS-telemetry dataset on a slow-life history migratory bird of prey – the European honey buzzard, tracked for multiple years after fledgling. Our aim was to assess how phenology, ranging behaviour and philopatry change as individuals gain experience. Individuals exhibited variability in the age at which they first returned to their breeding range, yet all survivors settled at sites proximate to their natal nests. In subsequent returns, immatures increased the time spent within the breeding range and narrowed down their ranging to smaller areas situated even closer to their natal nests. Our study unveils the complexity and protraction inherent in natal dispersal, emphasizing the importance of individual experience and migratory improvement for successful recruitment in long-lived species.

**HIGHLIGHTS:**

- We used a unique GPS telemetry dataset on honey buzzards to track their natal dispersal and recruitment in this long-lived avian migrant.
- All individuals gradually decreased their ranging areas, moved closer to natal sites, and increased the time spent on the breeding grounds to synchronize with adults and begin recruitment.
- Mortality during return migration was high and acted as a bottleneck.
- We found both processes to be complex and lengthy, with the overall duration depending on individual decisions regarding the timing of return migration.

## INTRODUCTION

Many animals depart from their natal habitat upon attaining independence. The phenomenon of permanent displacement from the natal area to the site of initial reproduction has garnered significant attention among ecologists and is extensively deliberated under the term “natal dispersal” (Greenwood and Harvey 1982). Natal dispersal strategy tend to vary greatly between and within species, and seem to emerge through a trade-off between establishing in familiar areas and habitats and conspecific attraction on the one hand, and avoiding intraspecies competition (Nilson 1989, Negro et al. 1997, Forero et al. 2002) and inbreeding (Alonso et al. 1998, Zedroesser et al. 2007, Ortego et al. 2008) on the other hand. Theprocess and outcome of natal dispersal not only influences the life history and fitness of individual organisms (Forero et al. 2002, Nevoux et al. 2013, Luna et al. 2020) but also shapes the genetic and social structure of populations (Ekman et al. 2001, Ortego et al. 2008, 2011).

Dispersal constitutes a movement process, yet in many investigations, it is often distilled to a single metric: dispersal distance (Benton and Bowler 2005). Undoubtedly, it is a much more intricate phenomenon and challenging to track, especially among long-lived species that mature late. In such species, dispersal unfolds as a multistep process. Between the initial departure from the natal site and settling in a new breeding ground for the first time, dispersing individuals undergo a wandering phase (Benton and Bowler 2005, Penteriani and Delgado 2009). During this stage, long-lived species, even upon reaching maturity, frequently linger for one or a few seasons, or even their entire lives as floaters, seeking a mate or suitable breeding territory (Penteriani and Delgado 2009), and this part of life is attributed to recruitment process. Recruitment can be defined as the establishment of new individuals in the breeding segment of a population (Reed et al. 2003), and the recruitment process is considered complete once an individual breeds for the first time. The age of first breeding is a key aspect of recruitment that significantly affects population demography (Gaillard et al. 1989, Stahl and Oli 2006). Along with clutch size and adult survival, it is one of the main components of birds’ life history strategies (Sæther and Bakke 2000). Species that mature early typically exhibit high fertility and low survival, characteristics of fast-life history species, while late maturation, low fertility, and high survival represent the other end of the life history continuum (Sæther 1988). However, the age of first breeding is also density-dependent, with younger individuals reproducing in low-density populations (Wyllie and Newton 1991), thus indicating the stage of population development.

In migratory species, wherein the period between leaving the natal site and initial breeding is preceded by one or more episodes of migration, dispersal is additionally complex. During this interim period, immature individuals may traverse thousands of kilometres to non-breeding grounds before returning to their breeding range. Inexperienced young bird migrants are often susceptible to deviations during migration due to navigational errors or wind drifts, resulting in their positions often shifting further and in a more random manner than those of resident species (Vickers et al. 2023). Consequently, migration profoundly influences the dispersal process, resulting in relatively longer dispersal distances (Paradis 1998) and relatively lower levels of philopatry (likelihood that individuals breed at or near their place of origin; Weatherhead and Forbes 1994) in species that migrate longer distances. At the same time, high philopatry (i.e. dispersal distances in the order of tens of meters to kilometers) tends to be widespread even among migrants (Förschler and Bairlein 2010, Ceresa et al. 2016) and even in species that spend long periods outside the breeding range before their first breeding attempt (Coulson 2016). In doing so, migratory birds, demonstrate remarkably long-term spatial memory and homing abilities, as well as a strong bias towards natal environmental cues during recruitment (Keiser 2003, Healy and Hurly 2004). And so the precise mechanisms through which early-life movements outside the breeding range influence the natal dispersal process remain obscure.

An additional challenge for migrants, especially those undertaking long-distance journeys, is that they spend relatively little time in their breeding range prior to recruitment. While immature individuals of resident species may spend the entire autumn, winter, and early spring exploring resources and assessing competitors’ distribution around potential breeding sites, migrants often have only a few weeks from arrival to recruit into the breeding population. In long-lived species with protracted offspring development and parental care, floaters, despite being physiologically mature, may struggle to secure a territory and breed within the short window following their initial arrival from wintering sites. In such species, the dispersal process is also protracted and typically encompasses the floater stage for at least one season. Importantly, individuals that do not improve their migration performance and arrival time might not even recruit into the breeding population (Sergio et al. 2017). All considered, this underscores the importance of better studying the link between migration timing and the natal dispersal and recruitment process.

The European honey buzzard (Pernis apivorus; therafter honey buzzard) serves as an example of a long-lived, long-distance migratory species with potentially intricate dispersal and recruitment strategies. Firstly, it is a specialist with a narrow diet, predominantly preying on Hymenoptera, especially wasps and their larvae (Gamauf 1999, van Manen et al. 2011, van Manen 2013) within tropical and temperate forests (van Manen et al. 2011, Howes et al. 2020a). This dietary specialization significantly influences its annual cycle and appears to constrain the suitable habitats where it can establish breeding populations. Secondly, migratory species of the honey buzzard genus undertake transcontinental migrations, covering several thousand kilometres between their temperate breeding and tropical non-breeding ranges (Higuchi et al. 2005, Strandberg et al. 2012), with juveniles and adults exhibiting a large ontogenetic shift in migratory behaviour (Kjellen et al 1992, Schmid 2000, Hake et al 2003, Vansteelant et al 2020). Thirdly, it is a secretive species, characterized by its brief residency in breeding areas, late spring arrival (during the full canopy season), and relatively low population density. Consequently, it has been inadequately studied, particularly in contrast to its relatively high conservation status (listed in Annex 1 of the Birds Directive).

Here, we present longitudinal movement data of young honey buzzards tracked from their natal areas in Finland for up to seven years after fledging, thus revealing their process of natal dispersal and recruitment (cf. Roberts and Law 2014). We specifically aimed to understand the interplay between maturity, natal dispersal, recruitment, and migratory behavior in this complex and secretive species. It is well established from migration counts, ringing and tracking data (reviewed in Vansteelant and Agostini 2021) that juvenile honey buzzards on their first outbound migration migrate later and over a broader front than adults, with relatively few juveniles migrating through traditional overland flyways (Kjellen et al 1992, Vansteelant et al 2020), and juveniles taking more time to complete their first migration (Hake et al. 2003). Juvenile behaviour thus seems to be consistent with an innate vector-based orientation strategy, making them highly susceptible to wind drift (Thorup et al. 2003). Indeed, a previous study based on the same individuals that we study here showed that stochastic wind influences greatly influenced the route choice of juveniles on their first outbound migration, as well as the longitude at which they settled in tropical Africa (Vansteelant et al. 2017). The remainder of the natal dispersal process, in particular the first return migration and breeding range movements of young honey buzzards, have not been tracked until now. We expect most honey buzzards to initiate their first return migration upon reaching sexual maturity in their 3^rd^ calendar year, and to home back near to their natal areas to recruit as breeders. We also expect to observe gradual changes in phenology and ranging behavior as the immatures mature, eventually resembling the patterns observed in breeding adults once individuals recruit.

## METHODS

### GPS-tracking dataset

As a part of ongoing work on honey buzzard migration and movement ecology (Vansteelant et al. 2017, Nourani et al. 2020, Howes et al. 2020, Brønnvik et al. 2022), young honey buzzards (n = 29) hatched in broods in southern Finland (latitude 61°1411–63°1211 N, longitude 21°1611–23°3111 E) were equipped with solar-powered Argos-GPS platform terminal transmitters (PTTs) (Microwave Telemetry Inc.) or Ecotone GSM-GPS-trackers (Ecotone) during 2011-2016 using body-loop harnesses made of Teflon ribbon. Tags’ weight (22–27 g) corresponded to ca. 3% of the birds’ body mass at deployment (908±83 g; avg ± s.d., n = 29). The amount of delivered GPS-fixes varied depending on tracker model and programming. On average, immature individuals collected 1180 (± 764 SD) GPS locations per individual per year. The sex of birds was determined from DNA as extracted from blood samples using the salt extraction method. For more details on the field protocol see Vansteelant et al. (2017). Additionally, a dataset on adult birds (n=9) was used for comparison of ranging areas on wintering grounds. The details of this dataset are described in Brønnvik et al.(2022).

### Ranging area

During consecutive wintering and breeding seasons, ranging areas were estimated to compare how the area of exploration changes with age and experience. The dataset comprised moderate-frequency GPS fixes, resulting in hundreds or occasionally dozens of fixes per season for some individuals. To compare ranging areas rather than estimate space use requirements, a geometric estimator – the 90% minimum convex polygon (mcp) – was employed. We considered alternative utilization distribution methods, but in case on non-territorial, wandering individuals of varying sample size and changing activity centres, utilization distribution methods overestimated the ranging area more than mcp.

Calculations were conducted using the adehabitatHR package (Calenge 2006) in R 4.1.1. Wintering ranging areas were calculated from the beginning of January until the end of March, when individuals had reached their wintering quaters and ceased moving long distances, until spring migration. Ranging areas upon returning to the breeding range were calculated from arrival until departure in the potential breeding range. By the latter we assumed southern Finland, where the tracked individuals originated and were expected to return, assuming the species is philopatric.

### Phenological dates

To track how phenology of migration changed with age, departure from non-breeding range, arrival to and departure from the breeding range were noted in the Julian calendar. Because individuals differed in the latitude of their ranging in Africa and this affects the distance and migration duration, the date of departure from non-breeding range was recorded as the date of passing the 13^th^ parallel (the approximate boundary of the desert zone). Arrival to the breeding range was determined as the first instance where an individual ceased moving northward by more than 50 km in the next 5 days. Departure from the breeding range was identified when an individual began moving southward by more than 50 km in the next 5 days.

### Philopatry

To assess if immature honey buzzards tend to return and potentially settle close to their natal sites, we calculated the mean and minimum distance at which honey buzzards resided from their natal site in each consecutive breeding period. To do this we used the ‘distance matrix’ tool in QGIS 3.22 for each breeding season.

### Mortality

Confirmed and probable deaths of tracked individuals, especially during their first autumn migration, wintering, and returning migration, were recorded to compare mortality among naïve honey buzzards. Suspected mortality was investigated on-site when indicated by sensors showing low temperatures, absence of movement over several days, or suspicious movements in urbanized habitats. Mortality due to drowning during migration across the Mediterranean and Black Sea was inferred from signal loss at the sea shore. Tag failure was suspected in some cases, especially in the fourth-seventh year of tracking. Mortality ratios were estimated based on confirmed cases, excluding suspected tag failures.

### Analyses

Maps were done in QGIS 3.22 (QGIS.org 2024). All the analyses were carried out in R 4.1.1 (R Core Team 2021). Due to sample size limitations, statistical approaches were constrained. Out of 29 individuals at the start, the sample size decreased to eleven individuals completing the first winter, six at the first successful return to the breeding range, four at the second, and three at the third. Wilcoxon tests were used to compare winter ranging areas between first and subsequent wintering episodes as well as to test sex differences in the year of first return to breeding range and date of arrival. Linear models were employed to test changes in dates of arrival and departure from breeding ranges (Julian date), duration of stay and size of ranging areas in breeding and wintering grounds (square km). Each of the above variables was tested in a separate model due to unequal and low sample sizes and explained by individual experience, measured by the number of returns to the breeding range or the number of wintering episodes in the case of winter ranging areas. Model residuals were assessed for normality with the Shapiro test and variables were log-transformed in the case of ranging areas.

## RESULTS

### Extended Immaturity Period in Wintering Range

Out of the 29 tracked juveniles, 22 (76%) successfully reached wintering sites in Africa (Tab. 1). None of the juveniles returned to the breeding range in their second calendar year. During their first winter, honey buzzards ranged over extensive areas in Central-Western Africa, with some individuals traversing distances exceeding 2000 km along longitudinal or latitudinal axes (Fig. 1A). Subsequently, some individuals reduced their ranging areas considerably during the next winter, while others maintained extensive ranges. However, the decrease in wintering area between the first and next winter was statistically insignificant (W = 57.5, p = 0.11), and no significant changes were observed in later life stages, although breeding adults tended to range over smaller areas (Fig. S1).

**Figure 1.**
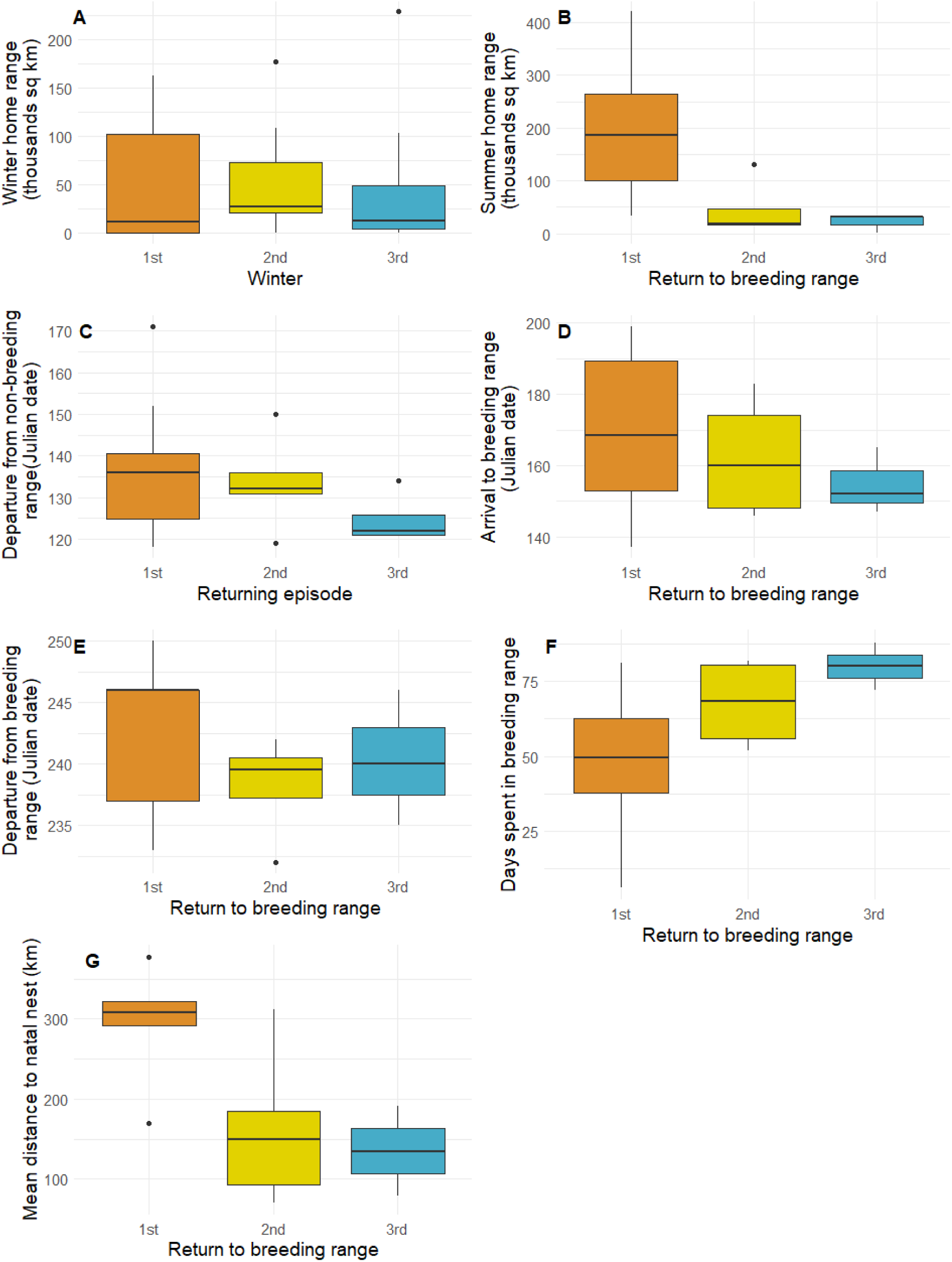
Differences in migration strategy and ranging behaviour with age (A) o or migratory experience (B-G) in GPS-tracked Honey Buzzards from Finland, followed from fledging. A - Size of ranging areas on wintering (January-March) and breeding (B) grounds, C - Date of departure from non-breeding range, D - Date of arrival to breeding range, E - Date of departure from breeding range (start of autumn migration), F - Duration of stay in the breeding range, G - Mean distance to natal nest during breeding season. Sample size decreased in time, from thirteen individuals followed through first winter, six at first return, four at second and three at third.

**Table 1.**
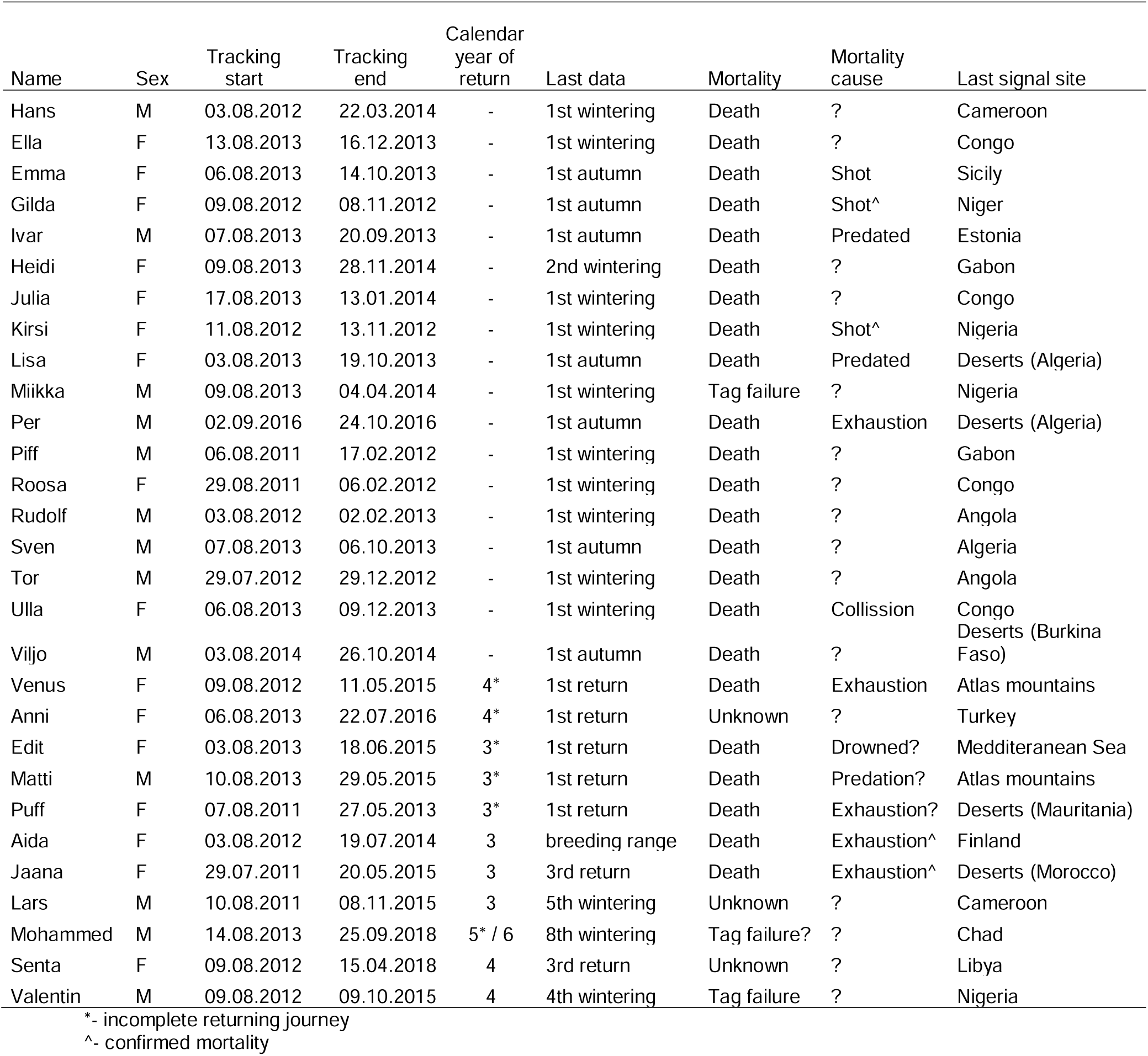
Characteristics of GPS-tracking dataset of twenty nine Honey Buzzards from Finland, followed from fledging.

In the 3^rd^ calendar year, six out of eleven tracked individuals attempted to return to the breeding range, with only three succeeding. The individuals that skipped the returning journey all survived to the next calendar year, and most of those (four out of five) attempted to return to the breeding grounds in the following year, in their 4^th^ calendar year. The last remaining individual, tried to return in his 5^th^ calendar year, but did not succeed in crossing the Mediterranean Sea and contined to wander in non-breeding part of the range (Fig. 2). Finally, this male successfully returned to his natal area in his 6^th^ calendar year. In total, only six individuals (21%) reached the breeding range between their 3^rd^ and 6^th^ calendar years. We did not confirm that establishing winter territories (represented by the size of winter ranging area) impacted the age of returning migration (W = 47, p = 0.76). Similarly, the change in winter ranging area between the first and second winters did not affect the decision to undertake the returning migration in the 3^rd^ calendar year (W = 10, p = 0.22).

**Figure 2.**
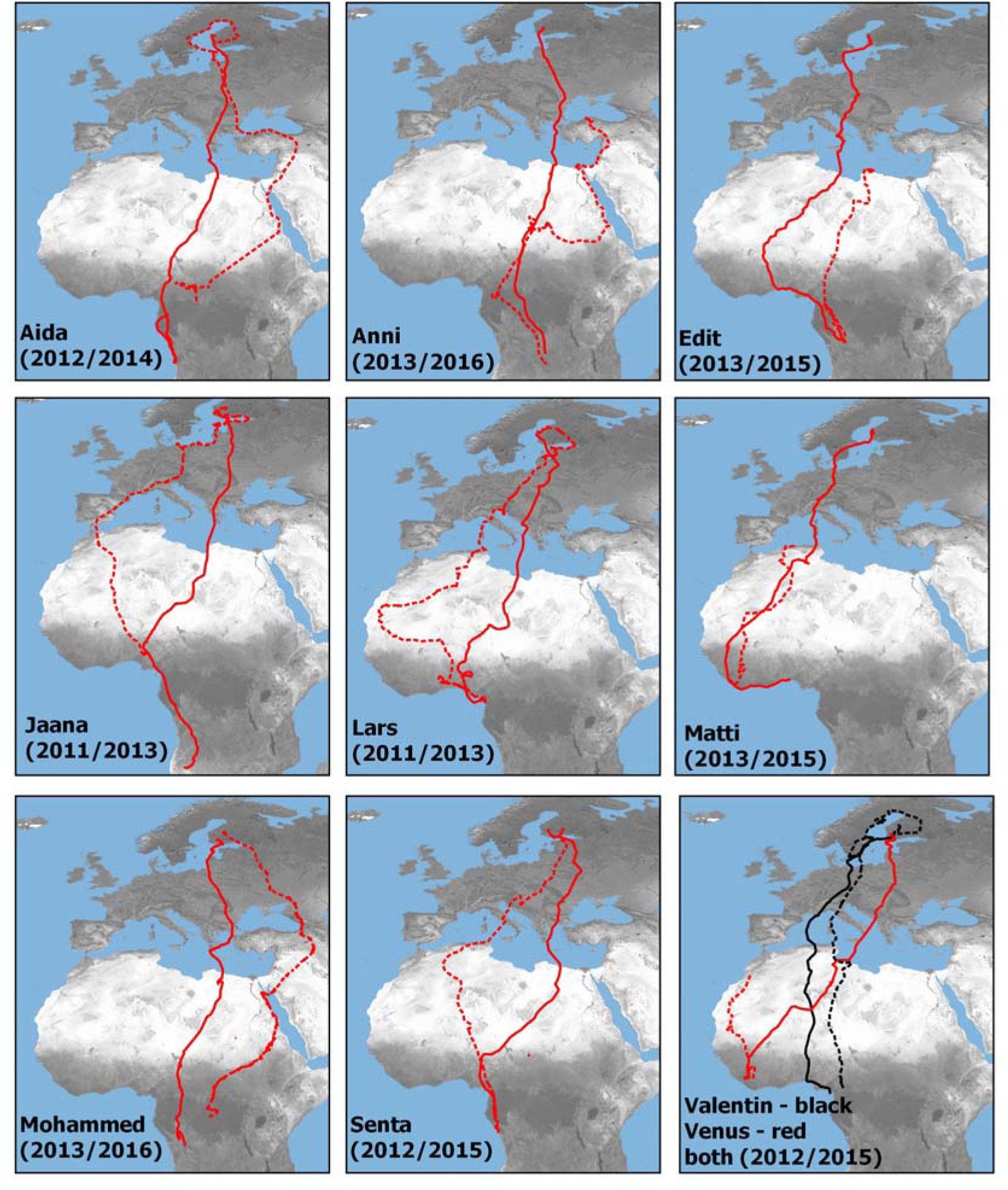
First autumn (continuous line) and first returning (dashed line) migrations of GPS-tracked Honey Buzzards from Finland. The names of individuals provided in each map correspond to the details given in Table 1. The year of the first autumn and returning migrations, respectively, are given in brackets.

### The First and Following Returns to the Breeding Range

All tracked individuals followed different paths during their first spring migration back to the breeding range when compared to their first autumn migration, with most paths not intersecting between autumn and spring (Fig. 2). Both clockwise and counterclockwise loop migrations occurred. Honey buzzards advanced their departure from the non-breeding range with each following return episode. The trend was visible (Fig. 1C), but only close to significant (Tab. 2). The date of arrival to the breeding range varied significantly between individuals especially during the first returning migration but tended to advance and narrow down slightly in subsequent returns (Fig. 1D, Tab. 2). The date of departure from the breeding range tended to be delayed across returns (Fig. 1E, Tab. 2). The mean departure date at first return was eight days earlier than the intial departure it their 1^st^ calendar year and about 2 weeks earlier at their second and third return. Immature honey buzzards spent more time in the breeding range with each subsequent return (Fig. 1F, Tab. 2). At first return, it was 22.2 days on average (range: 5 – 54), at second – 51.0 days (31 – 79) and at third – 63.7 days (56 – 84).

**Table 2.** Results of linear models explaining change in migration strategy and ranging behaviour with experience in GPS-tracked Honey Buzzards from Finland, followed from fledging (*p*-values are rounded to two significant digits, significant values are bolded).

| | Intercept | Beta<br>coefficient | Standard<br>error | $p$ | Adjusted $R^2$ |
| --- | --- | --- | --- | --- | --- |
| Departure from non-breeding range | 143.05 | -5.78 | 3.05 | 0.074 | 0.12 |
| Arrival to breeding range | 176.93 | -7.4 | 6.76 | 0.30 | 0.02 |
| Departure from breeding range | 242.89 | -1.3 | 2.12 | 0.55 | -0.06 |
| Duration of stay in the breeding range | 32.18 | 16.51 | 6.83 | **0.034** | 0.29 |
| Ranging area in breeding range (log) | 13.15 | -1.32 | 0.77 | **0.0064** | 0.46 |
| Mean distance to natal site | 376.23 | -92.53 | 31.54 | **0.014** | 0.39 |

### Recruitment and philopatry

The mean distance at which honey buzzards resided from their natal sites decreased significantly with each return (see Table 2, Figure 1G), averaging 296 km at the first return, 162 km at the second, and 135 km at the third (see Table S1). Similarly, the minimum distance to natal sites decreased with each return, ranging from 5 to 177 km at the first return (mean 53 km) to 0 to 174 km at the second return (mean 49 km). Five out of six returning individuals visited sites closer than 20 km from their natal sites, with four of them closer than 10 km. The area explored on the breeding ground by individuals tracked, at least until the second return, showed a significant reduction (Fig. 1B, Fig. 3). Following the first return, honey buzzards explored hundreds of square kilometres, whereas areas of only dozens of square kilometers were explored in the subsequent seasons. Some individuals confined their ranging to a portion of the area explored during the first return, while others moved to new, neighbouring areas that they had not explored before. Ranging areas showed significant overlap between the second and third return.

**Figure 3.**
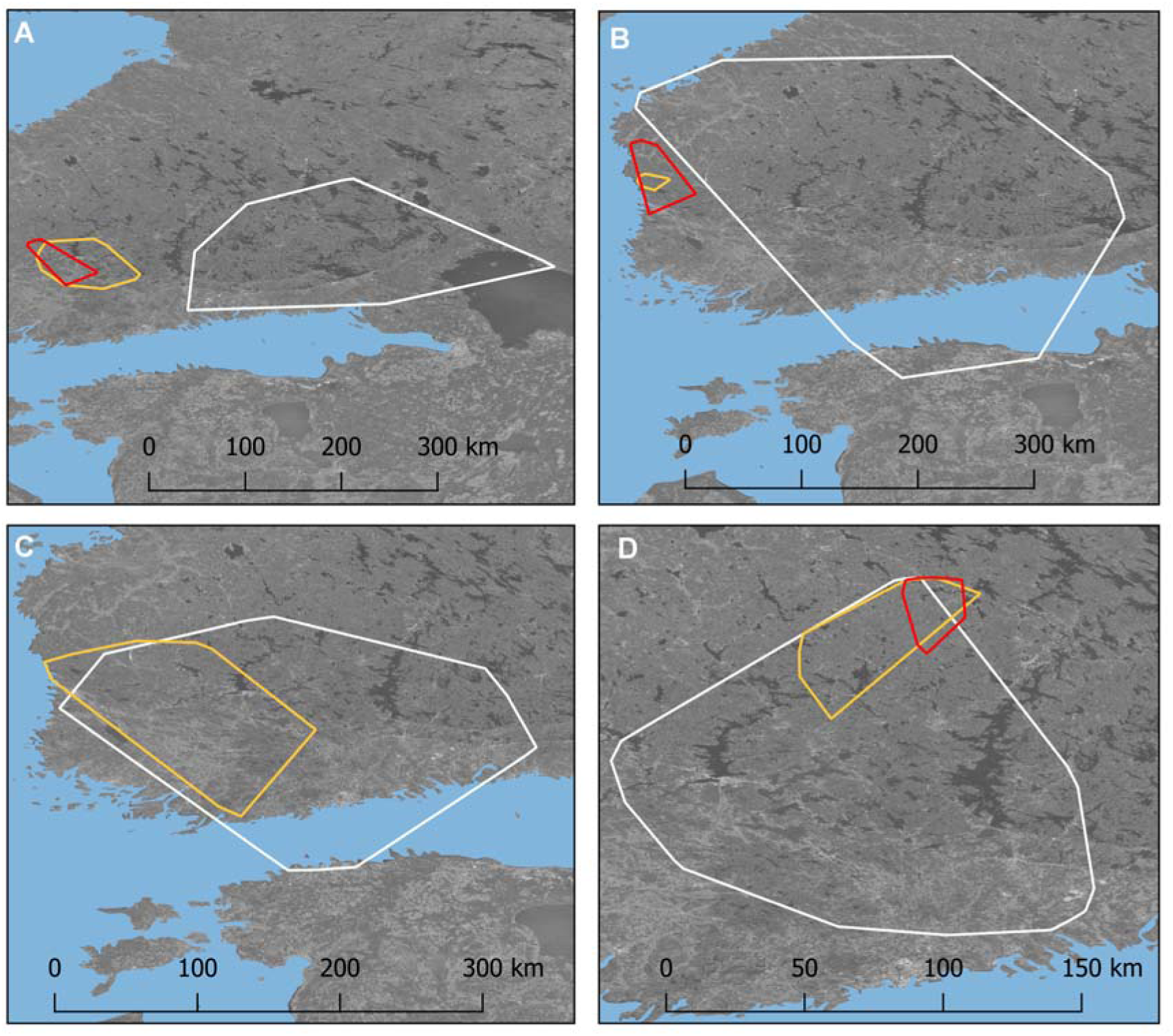
Change in summer ranging areas estimated with 90% minimum convex polygons of immature Honey Buzzards from Finland after first (white line), second (yellow line) and third return (red line). Plots A, B, C and D show ranging areas of Senta, Lars, Janna and Mohammed, respectively (see Tab. 1 for characteristics of individuals).

Two males were tracked long enough to locate their first nests and observe a central place foraging pattern around them (Tab. 1). However, we haven’t confirmed successful breeding. Lars built a nest in his 5^th^ calendar year, during his third return to the breeding range, 16 km from the natal site. Mohammed built his nest in his 6^th^ calendar year, during his first return to the breeding range, 189 km from the natal site. In his 7^th^ calendar year, he refurbished the same nest, but in the 8^th^, he probably built another one. Mating and breeding, however, were not confirmed in either of these males.

### Mortality Across the Long Dispersal Period

Twenty three of twenty nine tracked individuals (79%) were found dead. In the remaining six individuals (21%), the signal was lost, usually after 4-8 years of tracking (see Tab. 1 for individuals’ faith). In 4 cases (17%), we noted anthropogenic mortality, primarily due to shooting. Mortality on the first autumn migration reached 25% (7 out of 28 at this tage). All, except for one, managed to cross the Mediterranean Sea. Once they reached their wintering sites in the tropics, 32% of juveniles perished at wintering sites during the first wintering season and one more individual (3.5%) in the second season (altogether 10 out of 28). Finally, 39% of honey buzzards survived until their first returning migration. Out of 11 individuals that survived long enough to undertake the returning journey, as much as five (45%) perished trying to reach Europe. One individual turned back, so only less than 50% of individuals that survived to this stage were able to reach the breeding range in the first attempt and 55% in total. Once the honey buzzards successfully completed their dispersal cycle, only two mortality cases were noted. Overall, only 22% of initially tracked individuals managed to return to their breeding range, while only 15% lived up to their second return to the breeding range, when potentially the first breeding attempt was possible.

## DISCUSSION

Using GPS-GSM devices we were able to track the maturation and natal dispersal phase of honey buzzards from fledging to recruitment. Overall we reveal a remarkably protracted natal dispersal process, which sheds new light on the importance of individual experience and migratory improvements for successful recruitment in this long-lived species.

### Protracted immaturity period

Our study showed that the age of first breeding might vary between individuals, and that the honey buzzard breeds at a later age as compared to similar-sized raptor species, including resident species, and short-distance as well as long-distance migrants. While we could confirm nest-building by two GPS-tracked Finnish males in the 5^th^ and 6^th^ calendar years, breeding attempts could not be confirmed for other birds of similar age, and most individuals showed continuous changes in breeding range movements for as long as they were tracked. These results agree with studies on ringed British honey buzzards, which breed for the first time between the 4^th^ and 7^th^ calendar year (Roberts and Law 2014). The very late average age of first breeding is rather unexpected for a meso-predator like the honey buzzard. The similar-sized eurasian goshawk (Accipiter gentilis) can breed as early as the 2^nd^ calendar year, with as much as 42% of females of this resident species starting to breed that early (Krüger 2005). Short-distance migrant eurasian buzzards (Buteo buteo) can also breed early, but the average age of first breeding is 3.5 years (Davis and Davis 1992), and similarly short-distance migrant red kites (Milvus milvus) breed for the first time between the 3^rd^ and 8^th^ calendar year, with an average of 3.6 years (Newton et al. 1989). Long-distance migrant black kites (Milvus migrans) start breeding between the 2^nd^ and 8^th^ calendar year (Sergio et al. 2017). Overall, variability in the age of first breeding is expected in territorial, long-lived species that inhabit relatively stable environments. Prior to breeding, they need to establish their territories over high-quality sites, while non-territorial species inhabiting temporal habitats show much more limited variation in recruitment age (Charlesworth 1994, Aubry et al. 2009, Acker et al. 2018).

In the case of the honey buzzard, the late age of first breeding seems to be associated with a long delay of first return migrations, and an unexpectedly large variation in the age of returning individuals in their first returns. Individuals that return to the breeding range in their 3^rd^ calendar year could likely attempt breeding in their 4^th^ calendar year. By comparison, individuals like Mohammed (Tab. 1) that attempt a first return as late as the 5^th^ calendar year (or do not succeed to return before the 6^th^ calendar year) will recruit as breeders several years later. Such long delays of first return migrations, far beyond the age of sexual maturation and the attainment of adult plumage, were unexpected. The marked variation in the age of first return migrations, and the high survival rates of birds ‘oversummering’ in Africa at late age, suggests that honey buzzards face a trade-off between survival and attempting to return and recruit at early age. In this sense, it is striking that Mohamed, the bird to return at latest age, was one of the only birds for which attempted breeding could be confirmed in the field. It is possible that the age of first return is constrained by other factors like body condition or environmental factors, as has been posited for adult migrants that occasionally skip a northern summer by staying in sub-Saharan Africa (Sorensen et al 2017). However, the fact that at least half of all birds delay the first return at least two years, without engaging in failed or so-called ‘reverse’ migrations, suggest these between-individual differences reflect the viability of both ‘fast’ and ‘slow’ life-history strategies in this species. It is worth noting in this context that the longevity record in honey buzzards is 28 years, exceeding that of similarly sized raptors, being more similar to that of long-lived seabirds and vultures that also recruit at later age (Fransson et al 2023).

### Natal dispersal and philopatry

We followed the long process of natal dispersal in Honey Buzzards, although only single birds made it through to establish themselves as terrirtorial, nest-building individuals. Nevertheless, we were able to document the complexity of the whole multistep process preceeding breeding in a slow-life, transcontinental migrant species. All the tracked individuals found their way back to Finland and located themselves relatively close from their natal sites, which agreed with our expectations on the existance of philopatry in this species. The distance from natal nest and ranging areas decreased with age, while duration of the stay increased, which also met our expectations on individual improvements with age. This finding is in line with the results from the few studies documenting individual improvements by the means of GPS telemetry in slow-life history species. Consequently, in immature individuals of migratory Ferruginous hawks (Buteo regalis) returned significantly closer to their natal sites in their third spring (when they may already breed), than a year before (Watson et al. 2019). In partly migrating species, like the red kite, migration routes decresead in length along with birds age (Literák et al. 2022), also showing adjustment to adult-like behaviour.

Relatively little is known on how philopatric birds actually find their way back home. Do they repeat the autumn migration backwards up to familiar landscapes? Do they home by learned gradients in environmental cues (Thorup and Holland 2009, Kishkinev et al 2021) to return to the place they imprinted during the post-fledgling period? Our study adds insight to these uncertainities by showing that most individuals performed a loop migration. First return routes crossed or got close to first outbound routes in only two out of six individuals that completed their first return migration. Yet all birds managed to home back to Finland, even if some first had to return southwards after bypassing the Gulf of Bothnia through Sweden. To achieve such accurate navigation, birds may rely on a variety of mechanisms, such as bi-coordinate gradient maps, whereby they could infer their relative position by extrapolating from learned gradients in environmental cues (e.g. geomagnetic, olfactory). Imprinting on “home” takes place prior to abandoning natal sites (Sokolov 2000), as is evident from the fact that migratory birds that are displaced as eggs or chicks tend to return to the areas they were displaced to (Cade et al. 2000). Such imprinting takes time and might be one of the reasons why juveniles in many species, including honey buzzards, are migrating later than adults (Kjellen et al 1992, Hake et al. 2003, Vansteelant and Agostini 2021). Tracking data confirmed that juvenile Honey Buzzards were left by both adults at only 1 – 2.5 weeks after fledging (Zsiemer and Meyburg 2015). The extra time spent by juveniles around natal sites might therefore be used for further imprinting on the geographical location to which they will return via unknown routes, more than 18 months after they left.

### The process of recruitment to the breeding population

Tracking immature honey buzzards revealed that their recruitment to the breeding population greatly depended on their migration performance. Whereas relatively few unexperience juveniles died during their first autumn migration, survival was much lower during the returning migration. As a result, only about 22% of tracked individuals survived the returning migration bottleneck and were able to start the recruitment process.

We found that on their first return to the breeding range, immatures arrive much later than the adults, usually in mid-June, but even as late as mid-July. This is weeks after the breeding adults and certainly too late to breed. The time of arrival visibly advanced with each subsequent return, due to a combination of advanced departures from Africa and reduced migration time, which in turn was mostly due to reduced stop-over time in experienced migrants. Consequently the overall time spent in the breeding grounds extended, while departure timing from Europe in autumn tended to remain stable from the 2^nd^ autumn migration onwards. Already on the third return, honey buzzards spent on average 80 days on the breeding grounds, which is similar to adults (especially the ones that failed to breed). Therefore, the purpose of the first return in such long-lived migrant species seems to serve only for the exploration of the breeding grounds and possibly to initiate the annual routine and gain necessary experience in it.

Especially following the first return, honey buzzards clearly ranged over much greater areas in the breeding grounds than in following seasons. Wide-ranging behaviour is typical for floaters in many bird species (Tanferna et al. 2013, Eeden et al. 2017, Mirski and Anderwald 2023). Floaters are presumed to make such movements to obtain information on nest and foraging site distribution and quality, which can be inferred from conspecific performance too (see review by Lenda et al. 2012). The sharp decrease in ranging area and prolonged stay in the breeding grounds after the second return migration suggests that already at that time, honey buzzards might recruit to the breeding population, or at least have chosen the area of settlement. Indeed, two of the birds tracked here were witnessed to build nests after the 2^nd^ migration, in their 5^th^ and 6^th^ calendar year, respectively. Though nest-building does not yet equate to recruitment as an actual breeding individual, we based on our results find it is likely that most honey buzzards only engage in their first real breeding attempt after having experienced multiple migrations at 6-8 years of age. The exact age of first breeding in long-lived territorial species depends heavily on competitive skills (Becker and Bradley 2007)-which may include migratory performance (Sergio et al 2014, Flack et al 2024)- and habitat quality (van de Pol et al. 2007), with additional variation in our study species due to the variable age of first return.

## Conclusions

We found that both natal dispersal and recruitment in long-lived trans-continental migrants, such as honey buzzards, can be extremely long and complex processes. During the extended period of immaturity, individuals must familiarize themselves with wintering and breeding grounds, adjust their migration timing to match that of breeding adults, and navigate safely through the challenging spring conditions of the Mediterranean Sea. Finally, the age of recruitment in long-lived migrants can span greatly between individuals, largely due to the decision of when to return to the breeding grounds, which appeared to be a trade-off between survival and early recruitment.

## Ethics

Measuring and ringing of birds, was done as inherent of the normal ringing permit (permit 2604) as issued by the Finnish Museum of Natural History. All fieldwork requiring special permits (taking blood samples for DNA sexing, sampling and storing DNA-samples, attachment of PTTs/trackers) confirmed to five separate licences as issued by Finnish authorities (EPOELY/135/07.01.2013, ESAVI/2195/04.10.07/2014, PIRELY/49/07.01/2013, VARELY/73/07.01/2013, VARELY/215/2015).

## Data accessibility

The datasets generated and/or analyzed during the current study are available in the Movebank Data Repository, https://doi.org/10.5441/001/1.335 (Byholm et al., [2024]).

## Declaration of AI use

We have not used AI-assisted technologies in creating this article.

## Authors’ contributions

PM: conceptualization, data curation, formal analysis, investigation, methodology, visualization, writing—original draft, writing—review and editing; WV: conceptualization, methodology, writing— original draft, writing—review and editing; PB: conceptualization, data curation, funding acquisition, project administration, investigation, methodology, writing—original draft, writing—review and editing.

All authors gave final approval for publication and agreed to be held accountable for the work performed therein.

## Acknowledgements

We wish to thank Mikko Honkiniemi, Ari Rantamäki, Jari Valkama, Risto Vilén, Jouko Kivelä, Annika Rossi, Kari Palo, Hannu Vuoto, Daniel Burgas, Martti Peltola and Ismo Nousiainen for their help with tagging European honey buzzards in Finland.

