## Supplementary material for "Dispersal, maturation and recruitment in a long-lived, trans-continentally migrating bird": Online Supplement

### Supplemental Material

Table S1. Distance of immature Honey Buzzards from their natal sites, once returned to breeding range.

| Id | Sex | # return | Calendar year | Min (km) | Mean (km) | Comment |
|---|---|---|---|---|---|---|
| Aida | f | 1 | 3 | 177.4 | 295.3 | |
| Jaana | f | 1 | 3 | 27.3 | 291.0 | |
| Jaana | f | 2 | 4 | 7.0 | 149.3 | |
| Lars | m | 1 | 3 | 43.6 | 321.0 | |
| Lars | m | 2 | 4 | 12.5 | 92.8 | |
| Lars | m | 3 | 5 | 0.4 | 79.3 | |
| Senta | f | 1 | 4 | 47.2 | 321.6 | |
| Senta | f | 2 | 5 | 2.9 | 70.7 | |
| Valentin | m | 1 | 4 | 18.3 | 376.9 | |
| Valentin | m | 2 | 5 | 299.9 | 311.2 | Only 2 fixes |
| Mohammed | m | 1 | 6 | 5.0 | 169.2 | |
| Mohammed | m | 2 | 7 | 174.6 | 184.9 | |
| Mohammed | m | 3 | 8 | 174.9 | 190.8 | |

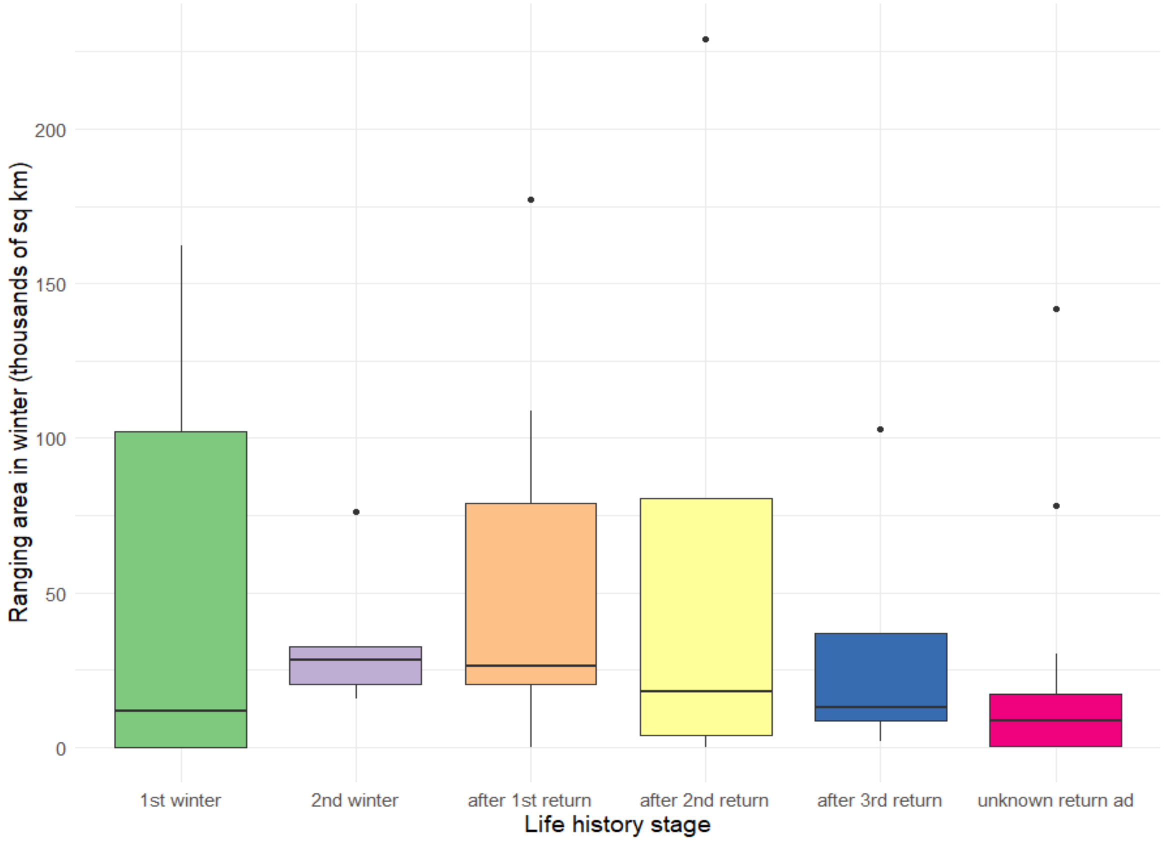


Figure S1. Winter ranging areas of Finnish Honey Buzzards in Africa across their life history. Ranging areas were estimated with 90% minimum convex polygon for the period of January-March. Note that some individuals spent more than two winters in Africa without returning to breeding range, while other overwintered only twice and started the migration cycle in their third calendar year. The last bar shows nine adults of unknown age, but caught already as breeders. In case of immatures, the sample size decreased in time, from thirteen individuals followed through first winter, six at first return, four at second and three at third.

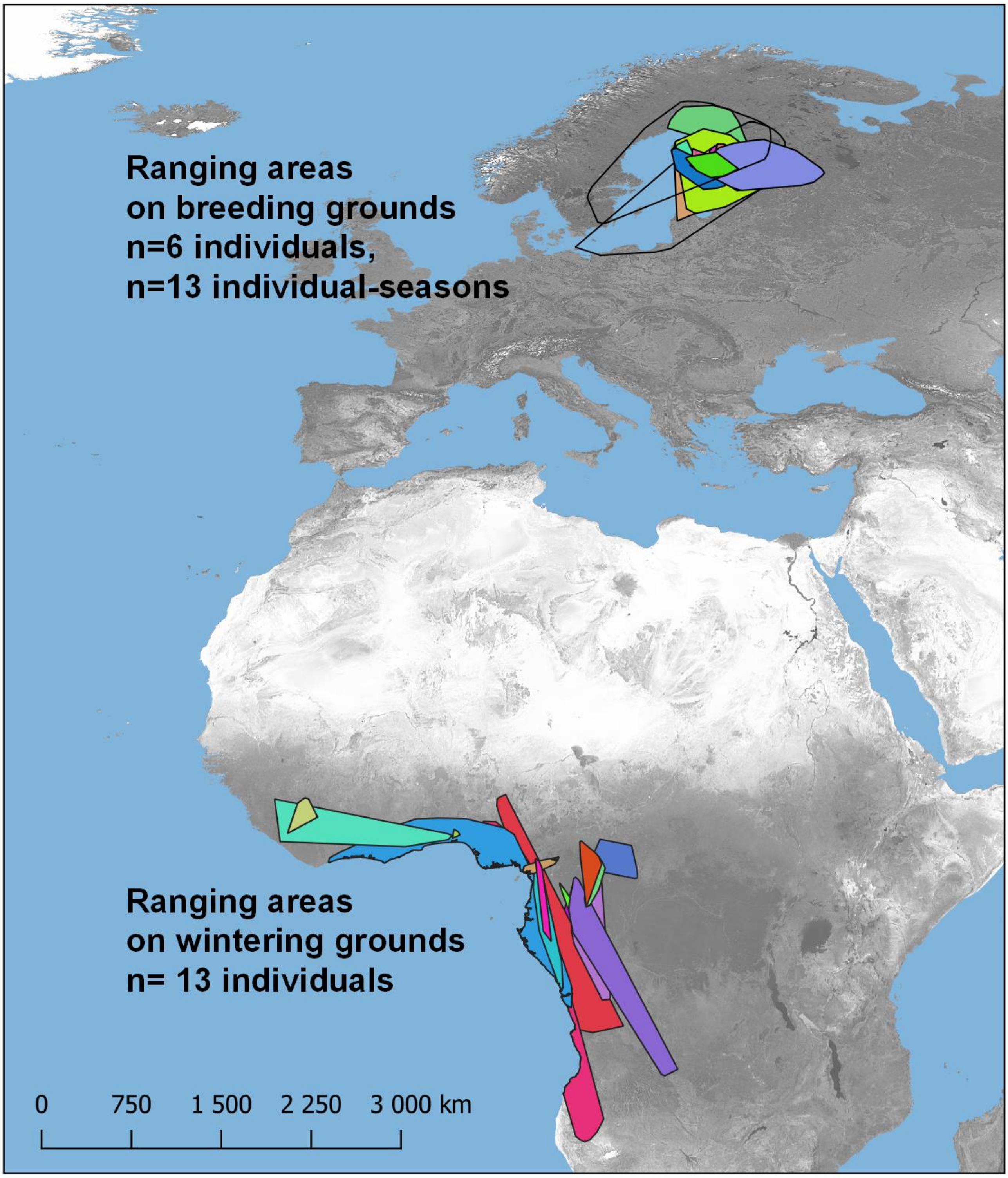


Figure S2. Ranging areas of immatrue Honey Buzzards on breeding and wintering grounds. (Each polygon fits to different year and individual)
